# Asymmetric horseshoe-like assembly of peroxisomal Yeast Oxalyl-CoA synthetase

**DOI:** 10.1101/2022.08.30.505785

**Authors:** Jérôme Bürgi, Pascal Lill, Evdokia-Anastasia Giannopoulou, Cy M. Jeffries, Grzegorz Chojnowski, Stefan Raunser, Christos Gatsogiannis, Matthias Wilmanns

## Abstract

Oxalyl-CoA synthetase from *Saccharomyces cerevisiae* is one of the most abundant peroxisomal protein in yeast and hence has become a model to study peroxisomal translocation. It contains a C-terminal Peroxisome Targeting Signal 1, which however is partly dispensable, suggesting additional receptor bindings sites. To unravel any additional features that may contribute to its capacity to be recognized as peroxisomal target, we determined its assembly and overall architecture by an integrated structural biology approach, including X-ray crystallography, single particle cryo-electron microscopy and small angle X-ray scattering. Surprisingly, it assembles into mixture of concentration-dependent dimers, tetramers and hexamers by dimer self-association. Hexameric particles form an unprecedented asymmetric horseshoe-like arrangement, which considerably differs from symmetric hexameric assembly found in many other protein structures. A single mutation within the self-association interface is sufficient to abolish any higher-level oligomerization, resulting in homogenous dimeric assembly. The small C-terminal domain of yeast Oxalyl-CoA synthetase is connected by a partly flexible hinge with the large N-terminal domain, which provides the sole basis for oligomeric assembly. Our data provide a basis to mechanistically study peroxisomal translocation of this target.

## INTRODUCTION

Oxalyl-CoA synthetase (OCS) is an acyl-CoA synthetase found in fungi and plants, but not in the vertebrate kingdom (Foster and Nakata 2014). OCS-catalyzed formation of oxalyl-CoA is crucial to prevent cellular toxicity due to accumulation of oxalic acid (Williams and Smith 1968). In contrast to plant oxalyl-CoA synthetases, the fungal enzymes comprise a C-terminal Peroxisome Targeting Signal 1 (PTS1) motif and hence are targets for translocation into peroxisomes. In yeast, OCS, also known as Pcs60, is one of the most abundant proteins in peroxisomes under conditions where peroxisome function is essential, such as growth in oleate (Blobel 1996).

OCS belongs to the acyl-CoA synthetase superfamily of adenylate forming enzymes (ANL) (Foster and Nakata 2014). Acyl-CoA synthetases are composed of two domains: a larger N-terminal domain (NTD) that comprises the active site, and a smaller C-terminal domain (CTD) that covers the NTD active site cleft upon substrate binding. Enzymes from the ANL family synthesize their products in a two-step reaction that requires a well characterized domain alternation mechanism (Gulick et al. 2003; 2009). The mechanism is associated with significant conformational changes, which involves a major rotation of the NTD/CTD arrangement.

To date, OCS structural characterization has remained limited to the non-peroxisomal *Arabidopsis (A.) thaliana* enzyme (Fan et al. 2016). Yeast OCS has become a model for PTS1 mediated cargo import, in which the PTS1 motif may become dispensable for peroxisomal receptor Pex5 binding, indicating a second non-PTS1 bindings site (Hagen et al. 2015). Based on these findings, we became curious about specific structural features of the peroxisomal enzyme, which may act favorably in promoting peroxisomal translocation. Surprisingly, by using X-ray crystallography, we found *Saccharomyces (S.) cerevisiae* OCS to be arranged in an asymmetric three-fold repeated layer of homodimers, which form a hexamer with an overall horseshoe shape. Using single particle cryo-electron microscopy (cryo-EM SPA) and Small Angle X-ray Scattering (SAXS), we confirmed that yeast OCS forms concentration-dependent single, double and triple-layered homo-dimeric assemblies with an overall configuration consistent with our X-ray crystallography data. A single mutation in OCS is sufficient to abolish the ability of OCS to form multiple layers of homodimers, which allowed us to characterize homogenous single layer OCS homo-dimeric species. In addition, we observed that *S. cerevisiae* OCS homodimerization is different from the *A. thaliana* enzyme and thus non-conserved. Finally, and again in contrast to *A. thaliana* OCS, the CTD in *S. cerevisiae* OCS is only in a loose arrangement with the NTD. Our data suggest that – unless these features have evolved for different functional reasons – they could play a role in promoting efficient *S. cerevisiae* OCS translocation into peroxisomes.

## RESULTS

### Yeast OCS forms an asymmetric three-layered hexameric assembly

To understand the structural arrangement of yeast OCS, we first determined the crystal structure of the enzyme at 2.9 Å resolution **(Figure 1, Supplementary Table S1).** Within each asymmetric unit of these crystals, we found two hexameric assemblies with an identical arrangement, composed of trimers of dimers **(Figure 1A, Table 1, Supplementary Figure S1)**.

**Figure 1:**
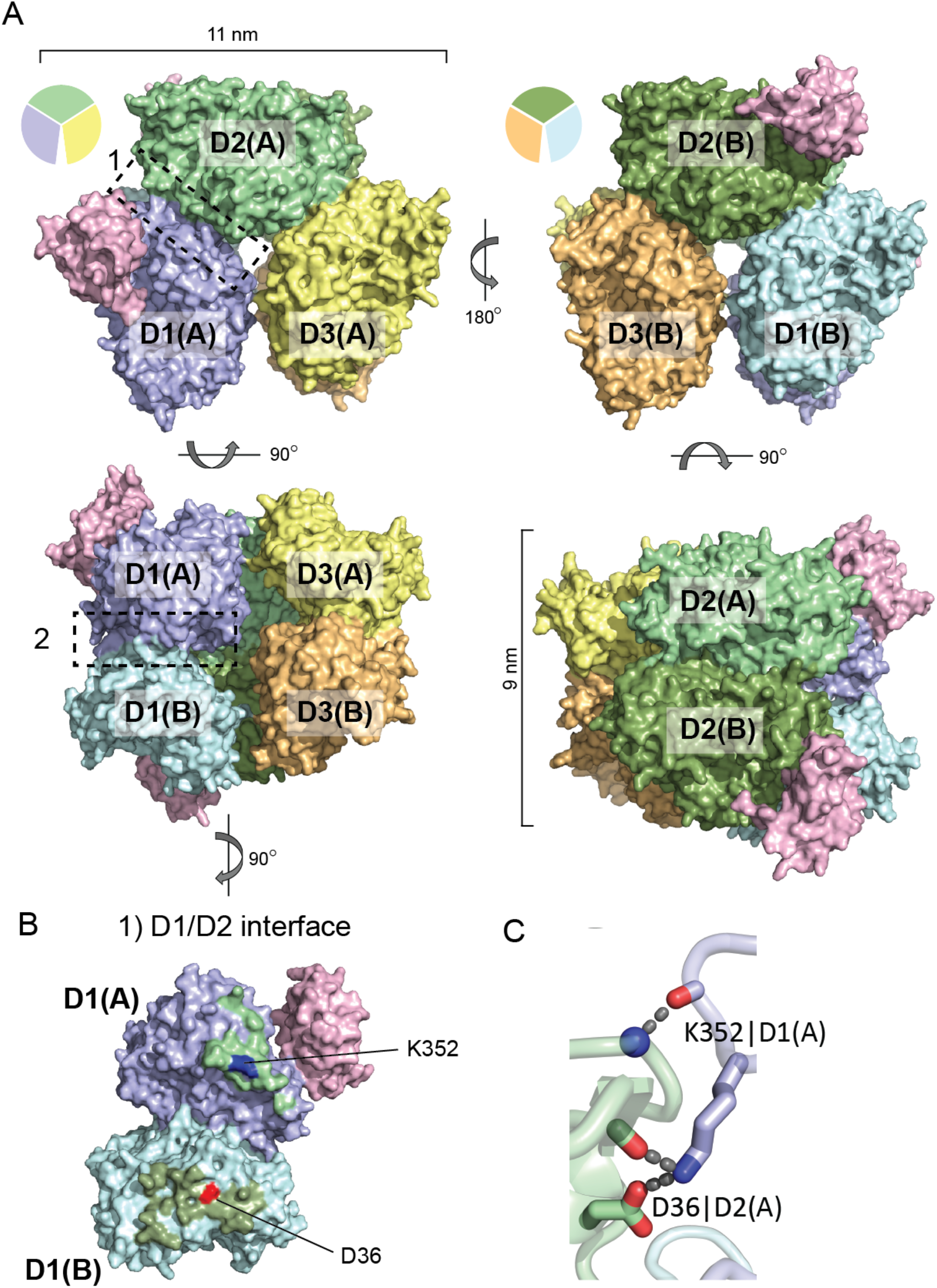
X-ray structure of hexameric OCS (wt) assembly. **(A)** Three OCS dimers associate into asymmetric hexameric assembly with a horseshoe-like overall shape. OCS dimer-1 protomers D1(A) and D1(B) are colored in marine blue and cyan, respectively; D2(A) and D2(B) in green and dark green, respectively; D3(A) and D3(B) in yellow and orange, respectively. Upper panel: view along a pseudo-threefold axis in two opposite orientations. As the protomers of the three OCS dimers are related by two-fold symmetry orthogonal to the pseudo-threefold axis, only the front layer of the three OCS dimers is visible. For the purpose of clarity, the visible OCS protomers are indicated by pie charts in the same colors and the angles, defining the relations between the three OCS dimers (*cf*. **Table 2**). Two CTDs out of six OCS protomers were included into the overall structure of the OCS hexamer-1, used in this illustration (in light pink). The OCS hexameric assemblies in the lower panel are rotated by 90 degrees around a horizontal axis with respect to the upper panel. Due to the overall horseshoe-like arrangement, a long deep crevice generated by the OCS hexameric assembly becomes visible (lower left panel). The overall dimensions of the OCS hexameric assembly are indicated as well. **(B)** Molecular details of OCS dimer/dimer assembly, exemplified for the D1/D2 interface indicated by box 1 in panel A. The D2/D3 interface is virtually identical **(Table 1)**. Colors for dimer D1 are as in panel A. Surface areas of dimer D1 that are in contact with dimer D2 are shown in the colors used for D2. Polar residues that contribute to specific dimer D1/D2 interface interactions are colored by their side chain charges (negative, red; positive, blue) and are labeled. The only sidechain specific polar interaction between D1 and D2 is generated by D26 and K352 and is two-fold repeated with opposite orientations, involving D1(A)/D2(A) and D1(B)/D2(B). The respective surface areas on D1(A, B) are boxed. **(C)** The D36-K352 interaction is supported by additional hydrogen bonds with the OCS main chain backbone (right panel).

**Table 1:** OCS oligomeric assembly.

| OCS (wt) |  |  |  |  |
| --- | --- | --- | --- | --- |
| Dimeric assembly | Rotation [deg.] | Interface protomers A/A [Å<sup>2</sup>] | Interface protomers B/B [Å<sup>2</sup>] | Total Interface [Å<sup>2</sup>] |
| D1 | 180 |  |  | 1009 |
| D2 | 180 |  |  | 1013 |
| D3 | 180 |  |  | 1020 |
| Hexameric assembly |  |  |  |  |
| D1/D2 | 113,5 | 398 | 397 | 795 |
| D2/D3 | 113.5 | 397 | 392 | 789 |
| D3-D1 | 133 | 24 | 15 | 39 |
| OCS (KD) |  |  |  |  |
| D1 | 180 |  |  | 1031 |

**Table 2:** OCS NTD/CTD arrangement flexibility.

| Subunit | CC <sup>a)</sup> | CTD sequence modelled <sup>b)</sup> | $\Delta(\Theta)$ NTD/CTD<br>$\Theta$ D1(A) = 0° <sup>c)</sup> |
| --- | --- | --- | --- |
| **OCS (wt) hexamer-1** |  |  |  |
| *D1(A)* | 0,83 | 443-533 | 0,0 |
| D1(B) | 0,59 |  |  |
| D2(A) | 0,36 |  |  |
| *D2(B)* | 0,76 | 443-519 | 10,0 |
| D3(A) | 0,17 |  |  |
| D3(B) | 0,31 |  |  |
| **OCS (wt) hexamer-2** |  |  |  |
| *D1(A)* | 0,21 |  |  |
| D1(B) | 0,41 |  |  |
| D2(A) | 0,70 | 443-467, 474-532 | 117,8 |
| *D2(B)* | 0,34 |  |  |
| D3(A) | 0,27 |  |  |
| D3(B) | 0,17 |  |  |
| **OCS (KD) dimer** |  |  |  |
| *D(A)* | 0,78 | 443-478, 489-507 | *n.d.* |
| D(B) | 0,53 |  |  |

<sup>a)</sup> Correlation coefficient (CC), as described in the Materials & Methods
<sup>b)</sup> Residue numbers as specified in P38137 (UNIPROT)
<sup>c)</sup> Calculated with DynDom (Lee, Razaz, and Hayward 2003)

While the electron density of all N-terminal domains (NTD) of the two OCS hexamers was well interpretable for the complete NTD sequence, the density accounting for the smaller CTD was less well defined to various degrees. We quantified the expected density for each CTD by map-model correlation analysis, revealing correlation coefficients (cc) between 0,15 and 0,83 **(Table 2).** As an analysis of contacts with symmetry-related OCS molecules did not reveal major differences, the varying levels of CTD density are a likely a consequence of inherent flexibility of the OCS NTD/CTD arrangement, in-line with previous findings of other members of the ANL family (Gulick et al. 2009). To this end, we used cc = 0,7 as a threshold for structural interpretation, to model two CTDs of the first OCS hexamer (chains D1(A) and D2(B)) and one CTD of the second OCS hexamer (chain D2(A)). Due to limited interpretable density, the CTD sequences that were modeled remained fragmented **(Table 2).** Modeling of these three CTDs allowed us to assigning OCS residues 1-429 to comprise the NTD, followed by a seven-residues flexible hinge segment (430-436), which connects to the CTD (residues 437-543).

The three OCS dimers found in both hexamers assemble into an unusual three-fold repeated stack, forming a horseshoe-like arrangement **(Figure 1A, Supplementary Figure S1).** This assembly is composed of two orthogonal layers where each of the three two-fold symmetry related OCS homodimers contribute one protomer to the first layer and the other protomer to the second layer, thus generating an asymmetric trimer of OCS dimers. The three-fold repeated OCS dimer layers are related by rotational angles between pairs of OCS dimers 1/2 and 2/3 of about 113 degrees each **(Figure 1A, Table 1)**, thus considerably deviating from three-fold symmetry. The two OCS layers with three protomers each are in congruent positions when looking along the pseudo-threefold axis. Within this OCS dimer layer arrangement, OCS dimers 1 and 2 as well as OCS dimers 2 and 3 interact via relatively small two-fold repeated interfaces of about 400 Å^2^ each (**Figure 1B, Table 1).** In these two virtually identical OCS dimer/dimer interfaces, we found a single side chain-specific salt bridge between D36 and K352 **(Figure 1C).** As result of these dimer/dimer arrangements involving OCS dimer 1 / dimer 2 and OCS dimer 2 / dimer 3, there is a deep and narrow groove between OCS dimers 1 and 3 (**Figure 1A, left panel**). The visible CTDs are not involved in any of these interfaces. In addition, each of the three OCS homodimers are formed through an elongated and two-fold symmetric interface of around 1100 Å^2^ between OCS NTDs, supported by an array of distinct specific interactions (**Figure 2A-B**). The overall dimensions of the NTD OCS hexameric arrangement are about 11 nm within each of the two trimeric OCS layers and 9 nm across the trimeric OCS layers **(Figure 1A).** When including the modeled CTDs the overall particle size increases to about 14 nm x 11 nm, respectively.

**Figure 2:**
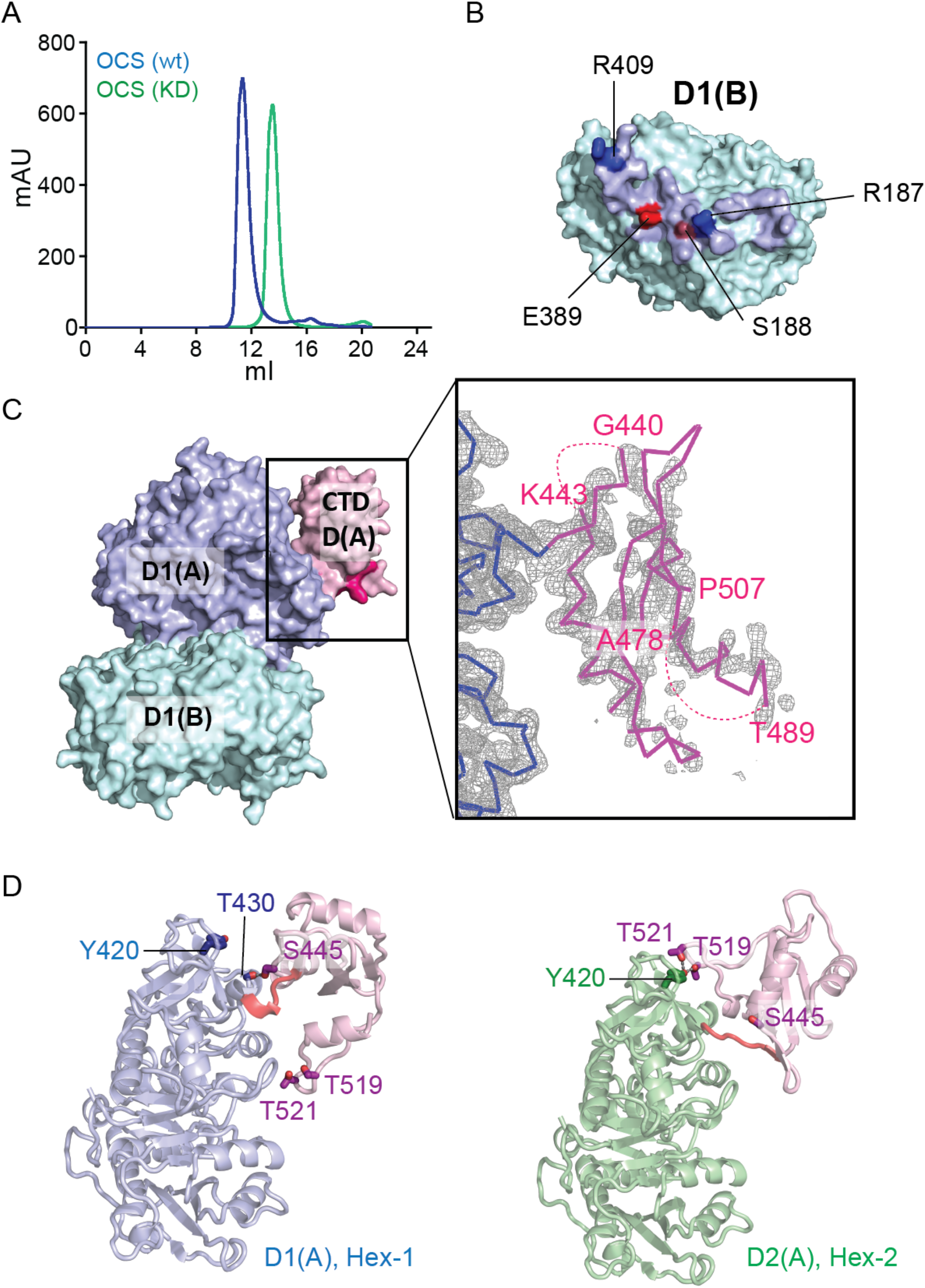
X-ray structure of dimeric OCS (KD) mutant assembly: **(A)** Size exclusion profiles of hexameric OCS (wt, blue) and dimeric K352D (KD, green). **(B)** Dimer interface of the KD variant, which is identical to the interfaces of the three dimers D1, D2 and D3 of the OCS (wt) hexameric assembly (*cf*. Figure 1). The colors are identical to those used for dimer D1 in the hexameric OCS (wt) assembly (Figure 1). Polar residues that contribute to specific dimer interface interactions are labeled. **(C)** Surface presentation of the dimeric OCS (KD) variant. The OCS CTD is visible in only in one of the two protomers of the OCS (KD) dimer (light pink). The NTD/CTD hinge region is colored in hot pink. Boxed inlet: composite *2Fo-Fc* OMIT map of the OCS (KD) D1(A) CTD in ribbon presentation. The boundaries of visible CTD fragments are labeled **(*cf*. Table 2). (D)** Molecular details of NTD/CTD interface of protomer D1(A) of the *wt* OCS hexamer-1 (left panel) and of protomer D2(A) of the *wt* OCS hexamer-2 (right panel). The CTD of D2(A) of the *wt* OCS hexamer-2 is rotated by 118 degrees with respect to the orientation of th CTD of D1(A) of the wt OCS hexamer-1 **(*cf*. Table 2**). This change is most significantly visualized by CTD residue positions T519 and T521, which are involved in specific interactions with the NTD in the second conformation only (right panel). The NTD/CTD arrangements found in protomer D2(B) of *wt* OCS hexamer-1 and D(A) of the OCS (KD) variant are similar to that of protomer D1(A) of the *wt* OCS hexamer-1 (left panel) (***cf*. Table 2, Supplementary Figure S1)**.

To investigate the OCS dimer module as a separate entity, we converted the positive charge of K352, which forms the salt bridge across the OCS dimer protomer interface, into a negative charge by mutating it to aspartate **(Figure 1C)**. As predicted, the OCS K352D (KD) variant lost its ability to form a hexameric assembly as seen by a delayed elution volume by size exclusion chromatography (**Figure 2A**). We then solved the X-ray crystal structure of the KD mutant to 2.5 Å resolution. In the crystal structure, the OCS (KD) variant forms a homodimer, which is identical to the one observed in the OCS (wild-type, wt) hexameric structure (**Figure 1A,** Figure **2B-C**). Similar to our observations in the hexameric OCS assembly there was only sufficient density for model building of a partially resolved CTD of one of the OCS protomers, supporting a semi-flexible arrangement of the NTD relative to the CTD in the crystal lattice (Figure **2C**). At this point we analyzed the nature of the NTD/CTD positions of all OCS protomers where it was possible to assign density to the CTD. While the arrangements of two OCS promoters of the wt hexameric assembly and of the OCS (KD) variant are identical, the CTD of the third OCS protomer with modeled CTD is rotated by 118 degrees when using the first NTD/CTD arrangement as reference **(Figure 2D, Table 2)**. The rotation is due to major changes of the dihedral angles of residues 433-435 within the NTD/CTD-connecting flexible hinge. Both arrangements are characterized by a scarce number of interactions between residues from the NTD and CTD **(Figure 2D).** While the first arrangement has a single hydrogen bond between main chain carbonyl group of T430 (NTD) and S445 (CTD) in common (**Figure 2D, left panel),** the second NTD/CTD arrangement is hold by a specific interaction between the tyrosine hydroxyl group of Y420 from the NTD and T519 and T521 from the CTD, close of the OCS C-terminus **(Figure 2D, right panel).** The latter two residues are not visible in some of the OCS protomers with the first NTD/CTD arrangement **(Table 2).** When comparing the two arrangements, for instance the positions of T519/T521 are different by about 3 nm on those superimposed protomers, where these residues are visible, well illustrating the scale of movement induced through this rotation.

### OCS dimeric layer self-assembly is concentration-dependent

We independently determined the structure of OCS by single particle cryo-EM (**Figure 3, Supplementary Figure S2, Supplementary Table S2**). After motion correction, the micrographs showed a homogeneous spread of globular particles **(Supplementary Figure S2A-B),** hinting towards OCS oligomers comprised of multiple copies of OCS homodimers. In addition to hexameric OCS accounting to 63% of all particles, we also detected populations of smaller particles with 33% associated with OCS tetramers and 4% with OCS dimers **(Figure 3A).** The 2D class averages accounting for hexameric assembly revealed a large complex with the same characteristic horseshoe-like assembly as observed in the crystal structure **(Figure 3A, right panel, Figure 1).**

**Figure 3:**
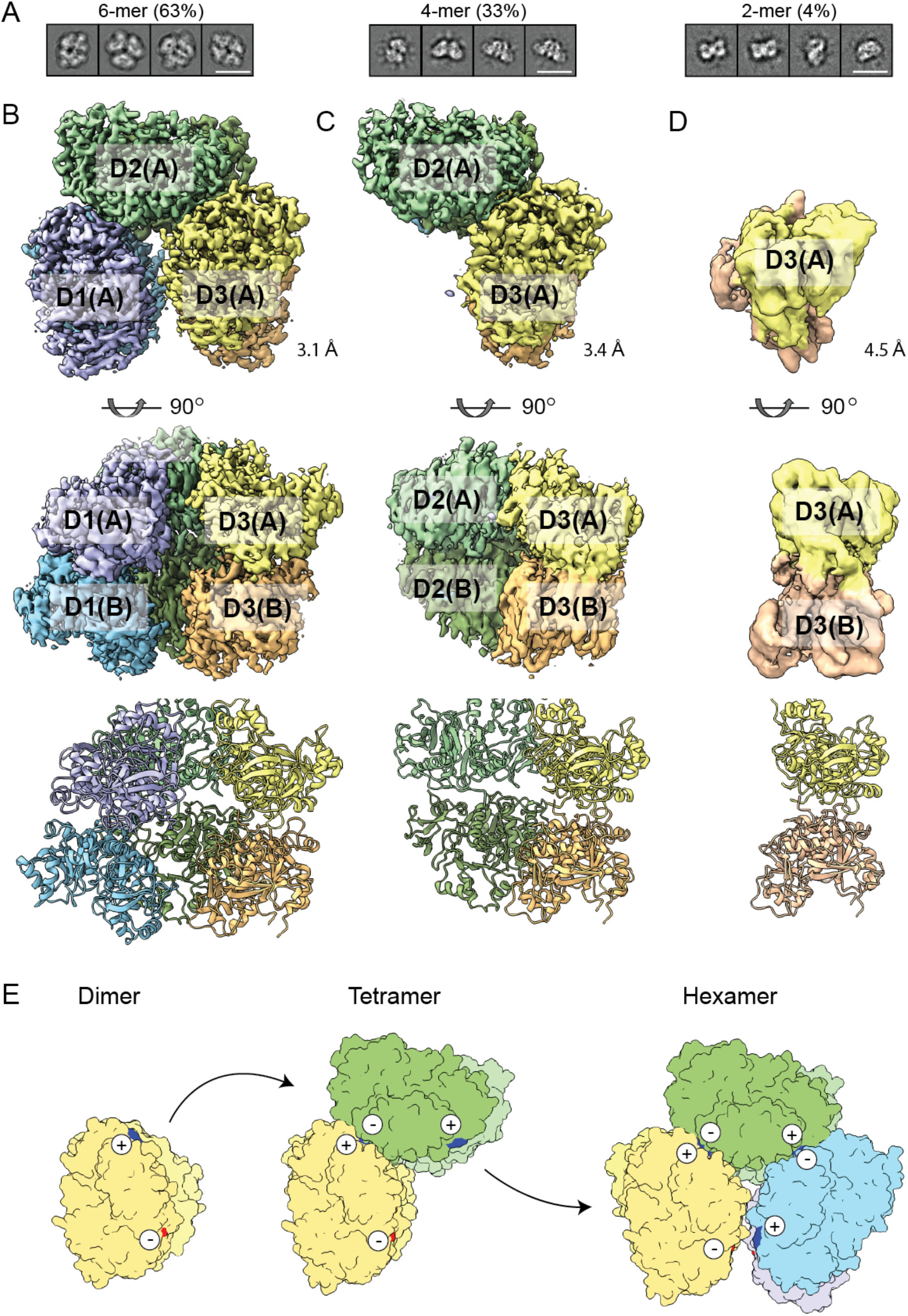
Cryo-EM structures of hexameric, tetrameric and dimeric OCS assemblies. **(A)** Representative 2D classes of OCS (wt) dimeric, tetrameric and hexameric assemblies, with the percentage of particles for each oligomeric assembly indicated. **(B)** Density maps of dimeric (left panel), tetrameric (central panel) and hexameric (right panel) OCS (wt) cryo-EM samples in two orientations, rotated by 90 degrees around a horizontal axis. The density map colors and labels are as in Figure 1. For comparison, structural cartoons of these assemblies are shown below. **(C)** Scheme indicating a plausible OCS dimer self-assembly path based on the distribution of cryo-EM particles and biophysical data (***cf*.** Figure 4). Dimeric OCS self-assembly requires the formation of a two-fold repeated salt bridge between neighboring OCS dimers (***cf*.** Figure 1). According to this scheme, OCS dimer self-assembly beyond the observed hexameric arrangement is not permissive, as this would lead to steric clashes.

For the hexameric OCS assembly, we obtained a 3D reconstruction at an average resolution of 3.1 Å (**Figure 3B, right panel, Supplementary Figure S3B**), which agrees well with the crystal structure. Hence, the complete crystallographic hexameric complex was first fitted into the cryo-EM density with a simple rigid body fitting and finally refined against the cryo-EM map. Unlike our observations in the OCS crystal structures, the cryo-EM map showed no density for any CTD, thus structural interpretation was limited to the analysis of the NTD arrangement (**Supplementary Figure S2B**). The root-mean-squares deviation (rmsd) between the refined cryo-EM and crystallographic model NTDs (based on Cα atoms of 2553 residues in total) was 1.26 Å, which indicate a high similarity between both models.

We further obtained cryo-EM structures of the tetrameric assembly and the OCS dimer, **(Figure 3B, left and central panel)**, which are however affected by anisotropic resolution due to preferred orientation of the respective particles. Therefore, we performed rigid body fitting of the available molecular model of the OCS dimers and did not attempt further model refinement. In summary, the distribution of these oligomeric states in solution indicates the tight OCS dimer as the basic building block. The tight dimers further oligomerize in a cyclic sequential manner via a single salt bridge towards the formation of the hexameric assembly **(Figure 3C)**.

Interestingly, we observed tetramers and hexamers only in cryo-EM samples and exclusively dimers during negative stain EM, that is typically performed at much lower concentrations (**Supplementary Figure S3A-B).** Chemical crosslinking before negative stain EM stabilized the hexameric horseshoe-like conformation **(Supplementary Figure S3C-D)**. These observations suggest that the protein might be prone to stepwise concentration-dependent self-association of OCS homodimers rather than concerted hexameric assembly.

To further characterize OCS dimer self-association, we analyzed OCS (wt) and the OCS (KD) variant at different concentrations by SAXS **(Supplementary Table S3**). OCS (wt) SAXS profiles show indeed a concentration-dependent change in the scattering intensities, as the sample was diluted from 7 to 0.5 mg/mL **(Figure 4A, Supplementary Figure S4A).** These changes correlate to a decrease in the apparent molecular weight (MW) and radius of gyration (*Rg*) of the particle population, consistent with mixtures of tetramers and hexamers, as observed in the cryo-EM samples **(Figure 4B, Supplementary Figure S4C-D)**. These observations for OCS (wt) contrast with those for OCS (KD) mutant, where the SAXS profiles and the corresponding MW and *Rg* as well as other structural parameters such as the Porod volume and real-space scattering-pair distance distribution are consistent with OCS dimers throughout the concentration series **(Figure 4C-D, Supplementary Figure S4B-D).** Indeed, by taking the X-ray crystal structures and subsequently including the mass and spatial disposition of both the NTD and CTD of each protomer in the OCS assemblies, the OCS (KD) mutant SAXS data is well described by the conformation of crystallographic dimer showing a χ^2^ fit = 1.0 (**Figure 4C)** whereas the fit to the OCS (wt) SAXS data requires the application of a volume-fraction weighted ratio of crystallographic hexamers and tetramers at each sample concentration resulting in χ^2^ fit range of 1.5 to 3.9 **(Figure 4A-B).**

**Figure 4:**
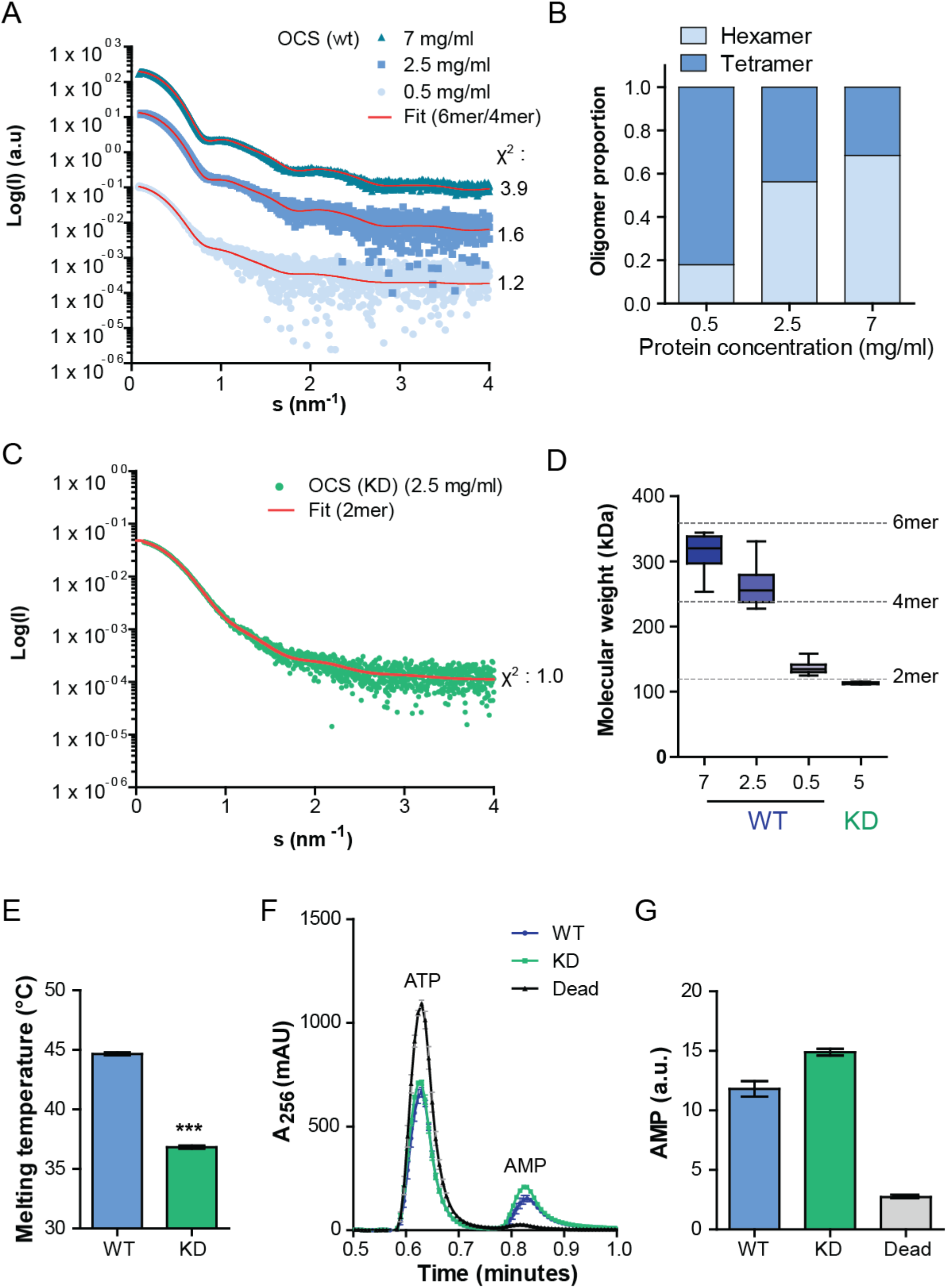
Biophysical and biochemical impact of concentration-dependent OCS assembly. **(A)** SAXS curves of OCS (wt) at different protein concentrations, with refined OCS tetramer/hexamer fit in red. Calculated χ^2^ values are indicated next to their respective curves. The SAXS curves are vertically offset for clarity. **(B)** Estimation of OCS (wt) tetrameric/hexameric assembly ratio according to protein concentration. OCS (wt) shows a shift from hexamer to tetramer with decreasing sample concentration. **(C)** SAXS curve of the OCS (KD) variant at 2.5 mg/ml with a complete OCS dimer model fitted. Calculated χ^2^ is indicated next to the curve. **(D)** Estimates of OCS (wt) and OCS (KD) MWs at different concentrations using SEC-MALLS. OCS (wt) shows a concentration-dependent change in MW, while OCS (KD) even at the highest concentration measured remains dimeric. **(E)** Melting temperatures of OCS (wt) and OCS (KD) measured by NanoDSF. OCS (KD) has a significantly lower melting temperature than OCS (wt). Measurements were carried out in triplicate. Student t-test ***, probability < 0.001. (F) Elution profile of 1µM OCS (wt), OCS (KD) and OCS (K523A) catalytic reaction after 1 minute with 100µM oxalate. ATP and AMP elute at distinct times and OCS (K523A) mutant show no formation of AMP. **(G)** Quantification of AMP products. Measurements were carried out in triplicate.

To confirm the concentration-dependent change in OCS oligomeric state, we also performed SEC-MALS experiments using three different OCS (wt) injection corresponding to concentrations of 7, 2.5 and 0.5 mg/mL sample. We observed a decrease in MW averages obtained for decreasing OCS (wt) sample injection concentrations, while even at the highest sample injection concentration tested for the OCS KD variant the MW average showed that the mutant MW corresponds to a dimer **(Figure 4D).**

### OCS oligomerization and thermostability

To investigate whether self-assembly of OCS dimers has an effect on thermostability, we determined the OCS (wt) and OCS (KD) variant unfolding temperatures using nanoscale Differential Scanning Fluorimetry (NanoDSF). While OCS (wt) unfolded at 44 °C, the dimeric OCS (KD) variant unfolded at 37 °C, corresponding to a reduction of 8 °C in melting temperature **(Figure 4E).** This implies that self-assembly of OCS dimers has a protective effect on OCS unfolding.

We finally tested whether lack of OCS dimer self-assembly impairs OCS catalytic activity, by monitoring ATP hydrolysis and AMP formation using reverse phase chromatography. As a negative control, we used an OCS K523A active site variant, which was expected to be catalytically inactive, in analogy to an equivalent mutant of *A.thaliana* OCS **(Supplementary Figure S5**) (Fan et al. 2016). As we did not detect any significant difference in the AMP levels generated (**Figure 4F-G**), we concluded that lack of OCS self-assembly has no impact on OCS catalytic activity.

## DISCUSSION

OCS is one of the most abundant proteins in yeast peroxisomes and comprises a PTS1 motif at its C-terminus (Blobel 1996). In contrast to the majority of other PTS1-containing peroxisomal targets, an OCS variant lacking the PTS1 is still capable of binding to the peroxisomal import receptor Pex5, indicating a second non-PTS1 receptor binding site (Hagen et al. 2015). To improve a mechanistic understanding of these findings, we compared the biophysical and structural properties of OCS from *S. cerevisiae* with the equivalent non-peroxisomal OCS from *A. thaliana*, presently the only other OCS with an experimental structure determined in a number of different ligand-bound and apo states (Fan et al. 2016).

The respective protomer structures can be superimposed with a root-mean-squares deviation (rmsd) of 1.8 Å for 418 matching residues of the NTD, which is equivalent to 96% of the complete NTD sequence **(Supplementary Figure S6A)**. In the resulting structure-based sequence alignment 40% of all residues are identical **(Supplementary Figure S5)**. Hence, it is not surprising that those parts of the structures of the two enzymes that are functionally important such as the active site are highly conserved.

At level of enzyme protomers, we have noticed that the NTD/CTD arrangement in the *A. thaliana* enzyme and the most populated NTD/CTD arrangement of *S. cerevisiae* OCS are identical **(Figure 3D, left panel; Supplementary Figure S6**). In both structures, the arrangement is mediated by an invariant serine (Ser445 in the *S. cerevisiae* sequence), following the flexible NTD/CTD-connecting hinge sequence segment, which belongs to the most conserved sequence segments in the overall OCS alignment **(Supplementary Figure S5).** In the *A. thaliana* apo enzyme (PDB entry 5ie0), however, there is a tartrate ion bound into the enzyme’s active site, which provides an additional specific interaction with K500 close to the C-terminus of the CTD (**Supplementary Figure S6B)**. In contrast, in all yeast OCS structures presented in this contribution, no ligand was found in the active site and the C-terminal segment of the CTD, which includes the equivalent lysine (K523), remains invisible. In summary, our data are suggestive that the found NTD/CTD arrangement is of functional importance irrespective a specific organism and absence or presence of a peroxisomal targeting signal. These finding are in-line with recent reviews on ANTs indicating that CTD flexibility may be a general ANT property (Gulick et al. 2009).

At the level of observed homo-dimeric assemblies, both enzymes form extensive assemblies by in part overlapping surfaces **(Supplementary Figure S5)**. Surprisingly, at the level of residue-specific interactions, the two interfaces are not conserved, thus leading to dimeric arrangements with the second protomer in distinct orientations **(Supplementary Figure S7).** Finally, the most striking difference is in the ability of *S. cerevisiae* OCS to form concentration-dependent asymmetric hexameric assemblies. The key residues permitting this assembly (D36, K352) are conserved in some fungi species, such as filamentous fungi *Chaetomium thermophilum* and *Neurospora crassa* and candida species (*Candida glabrata*), but not in fission yeast (*Schizosaccharomyces pombe and japonicus*), *Yarrowia lipolytica* and in plants (*A. thaliana*) (**Supplementary Figure S5)**. This may explain why such self-assembly property has not been observed for the *A. thaliana* enzyme. As the ability of forming higher order assemblies has been shown to be a property of the purified enzyme, it remains unknown whether such higher-order oligomerization is also formed under physiological conditions and possibly promotes translocation into yeast peroxisomes. Irrespective of its possible functional impact, it remains an unusual structural feature, which to the best of our knowledge has never been observed in any other protein hexameric assembly (Maksay and Marsh 2017; Marsh and Teichmann 2015). Although an increasing number of deviations to the original notion by Goodsell “*Asymmetric chitooligomers are virtually unknown”* (Goodsell and Olson 2000) have been found, the asymmetric self-association of OCS dimers remains a fascinating property of the *S. cerevisiae* enzyme. Taking our data together, whether and to what extent this collection of distinct properties provides any advantages to promote peroxisomal translocation remains subject of future research.

## MATERIAL AND METHODS

### Protein purification

Full-length *S. cerevisiae* OCS was cloned in a petM14 vector. OCS was expressed in autoinduction medium (Studier 2005), for 5 hours at 37°C and 26 hours at 20°C. Cells were harvested, resuspended in lysis buffer (50 mM Hepes (pH 7.5), 300 mM NaCl, 20 mM imidazole, protease inhibitor (Roche), DNAse (Sigma) and lysozyme (Sigma), homogenized for 1 hour at 4°C and lysed by sonication. Lysate was then cleared by centrifugation and the supernatant filtered at 0.45 µm and loaded onto Ni-NTA resin. Bound proteins were washed with 50 mM Hepes (pH 7.5), 500 mM NaCl, 20 mM imidazole. Protein was eluted with 50 mM Hepes (pH 7.5), 150 mM NaCl, 250 mM imidazole. The eluate was then dialyzed against Hepes (pH 7.5), 150 mM NaCl, 0.5 mM TCEP and simultaneously digested with 1 mg TEV-protease. Undigested protein and TEV protease were removed by a second Ni-NTA step and flowthrough containing OCS was concentrated to 5 ml for subsequent gel filtration (Hiload 16/60 Superdex 200 pg, GE healthcare). Relevant fractions were pooled together, and the protein was concentrated, flash frozen in liquid nitrogen and stored at −80 °C.

### Crystallization and X-ray structure determination

OCS (wt) was crystallized by vapor diffusing using conditions from the PEG I crystallization suite (Molecular Dimensions). OCS (wt) at a concentration of 6 mg/ml was crystallized in 0.2 M LiCl, 0.1M Hepes (pH 8.0), 15 % w/v PEG-3350. The OCS KD variant was crystallized in 0.15 M ammonium sulfate, 0.1 M MES (pH 6.0), 15% (w/v) PEG-4000. Crystals were soaked in cryo-solution, containing the crystallization mother liquor supplemented with 25% [v/v] ethylene glycol. Crystals were then mounted on cryo-loops (Hampton Research), and flash-cooled in liquid nitrogen. X-ray diffraction data of OCS (wt) and the OCS (KD) variant were collected on the synchrotron radiation beamlines MASSIF-1/ID30A-1 (ESRF, Grenoble) and EMBL P13 at PETRA III (EMBL/DESY, Hamburg), respectively. Data were processed using XDS (Kabsch 2010).

The OCS (wt) structure was solved by molecular replacement (MR) with PHASER (McCoy et al. 2007) using 4-Coumaroyl-CoA Ligase from *A. thaliana* (3TSY) as a search model. The initial MR solution was rebuilt interactively in COOT (Emsley et al. 2010). Each asymmetric unit contains 12 OCS protomers, grouped into two OCS hexamers, which are virtually identical in the OCS NTD arrangement (GESAMT (Krissinel 2012) superposes 2,532 C-alpha atoms of the two hexamers NTD domains with rmsd of 0.525 Å). To simplify building CTD fragments that were relatively poorly resolved in the map, we used AlphaFold2 version 2.1.1 predicted structure (plddt > 90) (Jumper et al. 2021). The predicted CTD structure was rigid body fitted to the map and real space refined in COOT with self-restraints generated at 5Å cut-off for all the chains. After subsequent refinement in REFMAC (Murshudov et al. 2011) version 5.8.0267 with NCS restrains and automatically defined TLS groups, CTDs with map cc’s < 0.7 (CC_mask calculated using phenix.map_model_cc (Liebschner et al. 2019)) were completely removed from the model. From remaining CTDs, residues without interpretable 2FoFc density at 10 level were removed. The resulting model contained two OCS CTDs for first hexamer and one CTD for second hexamer (***cf*. Table 2).** The final model was refined to R/Rfree of 0.23/0.25 at 2.87Å resolution.

The OCS (KD) mutant structure was solved by MR with PHASER using the OCS (wt) NTD (residues 1-431) as a search model. The model was completed with AlphaFold2 predicted CTD structure and refined as described above. For the mutant structure we built a fragmented CTD for one out of two OCS chains in the asymmetric unit (***cf*. Table 2).**. The final model refined to R/Rfree of 0.21/0.25 at 2.45Å resolution. Further details are summarized in **Supplementary Table S1.** The X-ray structures of OCS (wt) and OCS (KD) have been deposited in the Protein Data Bank with the following codes, respectively: 8AFF, 8AFG.

OCS interfaces were analyzed using the program PISA (Krissinel and Henrick 2007). Angles between OCS (wt) dimers were calculated using the PSICO (https://github.com/speleo3/pymol-psico) plugin in Pymol (DeLano 2020). NTD/CTD arrangements were analyzed with the on-line version of DynDom (Hayward and Lee 2002; Veevers and Hayward 2019).

### Negative Stain EM

Four µl of OCS sample in TBS Buffer (50 mM Tris pH 8.0, 150 mM NaCl) was applied for two minutes on a freshly glow discharged carbon coated copper-grid (Agar scientific; G2400C) at room temperature. The sample was blotted with Whatman paper (No 5) and subsequently the grid was washed two times using 10µl using TBS buffer. A 10µl drop of freshly prepared 0.75% uranyl formate solution was then applied on the grid for 30 seconds and the specimen was air-dried. Images were recorded with a JEM-1400 (JEOL) or a FEI Tecnai G2 Spirit (FEI) both equipped with LaB6 cathode and a 4K CMOS detector F416 (TVIPS). Single particles were selected using cryolo (Wagner et al. 2019). Reference-free stable class-averages were computed using the ISAC approach of the SPHIRE software package (Moriya et al. 2017).

### Crosslinking

Crosslinking with glutaraldehyde was performed in 50 mM Hepes (pH 8.0), 150 mM NaCl. Reaction mixtures with an amount of 0.003 mg/ml OCS were treated with 0.05% crosslinking reagent for two minutes and afterwards quenched by adding Tris (pH 8.0) to a final concentration of 100 mM. Prior to negative stain EM imaging, samples were purified via SEC using a Superose 6 Increase 5/150 GL.

### Cryo-EM structure determination

For cryo-EM, four µl of sample at 0.3 mg/ml concentration was applied to freshly glow discharged holey carbon Quantifoil 2/1 grids (#EMS300-Cu). The sample was immediately blotted and plunged into liquid ethane cooled by liquid nitrogen using a Gatan Cryo-plunge 3 device at 95% humidity. The blotting time was 2.5 seconds at 25 C (gentle blot). A cryo-EM dataset was collected with a Cs-corrected Titan Krios EM (ThermoFisher) equipped with a K2 direct electron detector (Gatan) and a quantum energy filter (Gatan). Movies were recorded in counting mode using the automated acquisition program EPU (ThermoFisher) at a magnification of 105,000x corresponding to a pixel size of 1.09 Å. 5594 movies were acquired in a defocus range of −0.7 to −2.2 µm. Each movie comprised 60 frames acquired over 12 seconds with a total dose of ∼63 e^-^/Å^2^.

Monitoring of image collection as well as image pre-processing was performed using the TranSPHIRE software package (Stabrin et al. 2020). Motion correction was thereby performed on the fly using MotionCor2 (Zheng et al. 2017). The contrast transfer function was estimated using gCTF (Zhang 2016). Ten representative images were then manually picked using the crYOLO-box manager tool, to train crYOLO(Wagner et al. 2019) for further automated particle picking. A total of four million particles were finally selected and extracted using a box size of 182 pixels. SPHIRE (1.0,1.2,1.3) (Wagner et al. 2019) was used for further single particle analysis steps. Multiple reference-free 2D classification rounds were then carried out, using the stable alignment and cluster approach (ISAC) (Yang et al. 2012) in SPHIRE **(Supplementary Figure S5**). After each round of 2D classification, we selected classes containing intact OCS populations, comprising OCS dimers, tetramers and hexamers. The resulting class averages were selected and subsequently used for VIPER, to generate a low-resolution initial model. The respective class members for each oligomerization state were then subjected to 3D refinement in SPHIRE (Meridien) with C1 (dimeric and tetrameric particles) or C2 (hexameric particles) symmetry, imposed using the VIPER reconstruction as initial reference, respectively. The resulting projection parameters were used to re-center and re-extract the particles. The re-centered particles were again subjected to 3D refinements in SPHIRE (Meridien). Particle “polishing” was then performed using RELION (Zivanov, Nakane, and Scheres 2019). Final refinements were then performed in SPHIRE using the polished particles of each individual population, respectively. Fourier shell correlation analysis, 3D masking, sharpening and low-pass filtering were automatically performed using the PostRefiner tool of SPHIRE. Further details are given in **Supplementary Table S2.**

Cryo-EM maps and models were visualized using Chimera (Pettersen et al. 2004) and ChimeraX (Pettersen et al. 2021). Map segmentation was performed using the module Segment Map of the Chimera software package. Structure modeling of OCS was performed using COOT (Emsley et al. 2010). Real space refinement of the crystal structure in the EM density map was performed in PHENIX (Adams et al. 2011).

The resulting model coordinates are available in the Protein Data Bank wth the PDB ID 8ATD. The cryo-EM structures have been deposited into the Electron Microscopy Data Bank with the ID EMD-15646.

### Small angle X-ray scattering analysis

Synchrotron SAXS data (*I*(*s*) vs *s*, where *s* = 4πsin*θ*/λ, 2θ is the scattering angle and λ the X-ray wavelength, 0.124 nm) were measured from dialyzed solutions of OCS (wt) and OCS (KD) in 50 mM Hepes (pH 7,5), 150 mM NaCl, 0.5 mM TCEP at the EMBL P12 beam line at DESY (Blanchet et al. 2015) (Hamburg, Germany) using a Pilatus 6M detector at a sample-detector distance of 3 m. OCS (wt) and OCS (KD) protein solutes at 7 or 5, 2.5 and 0.5 mg/ml concentrations were measured at 10° C. A continuous flow 1 mm cell capillary was used to limit radiation damage. A total of 30 successive 0.1 second frames were collected for both the samples and the corresponding matched buffer. The process of 2D data reduction to generate 1D buffer-subtracted profiles was performed using the SASFLOW pipeline (Daniel Franke, Kikhney, and Svergun 2012).

Data were analyzed using modules of the ATSAS 3.0.1 package (D. Franke et al. 2017). CRYSOL (Svergun, Barberato, and Koch 1995) was used to calculate the scattering profiles from the atomic coordinates of the OCS (wt) and OCS (KD) X-ray structures. CTDs were added to OCS protomers without CTDs modeled in the respective X-ray structures to create complete OCS model and fit them with the respective experimental SAXS data.

To characterize mixtures of the different oligomeric OCS species, the program OLIGOMER (Konarev et al. 2003) was used. It fits the observed experimental data by a weighted combination of the calculated model scattering curves (form factors) from different quaternary structures. The experimental SAXS data and models are deposited in SASBDB (Kikhney et al. 2020) (https://www.sasbdb.org/) with the accession codes SASDPV3, SASDPW3, SASDPX3, SASDPY3. Further details are presented in **Supplementary Table S3.**

### Size-exclusion Multi Angle Light Scattering

Protein mass measurements were performed on an Agilent HPLC system connected to a MiniDAWN® TREOS® multi-angle laser light scattering detector (MALS), with an Optilab T-rEX (RI) refractometer (Wyatt, Germany). 50 µl of sample at 10, 5, 1 mg/ml for Ocs (WT) or 5 mg/ml for Ocs (KD) were loaded onto a Superdex 200 Increase 5/150 GL (Cytiva) and the MALLS and RI measurements were performed at 25 °C, where the refractive index increment (dn/dc) of the protein samples was set to 0.185 mL/g. The data were processed using Wyatt ASTRA 7.0.1 software (Wyatt).

### Nano differential scanning fluorimetry

Melting temperature were measured by NanoDSF using a Nanotemper Prometheus NT.48. Tryptophan intrinsic fluorescence emission was recorded at 330 and 350 nm with 30% excitation power during thermal ramping from 20 °C to 90°C at a ramping velocity of 1°C per minute. A volume of 10μl OCS in 50 mM Hepes (pH 7.5), 150 mM NaCl, 0.5 mM TCEP and at a concentration of 1 mg/ml were loaded in NanoDSF grade standard capillaries. Samples were measured in triplicates and the calculated melting temperatures averaged. Melting temperature were calculated by the software PR. ThermControl (Nanotemper).

### Enzymatic activity assay

Enzymatic activity was measured by AMP production at 254 nm after 60 seconds reaction at 25 °C. The sample volume was 40 μl, containing 50 mM Hepes (pH 7.5), 150 mM NaCl, 2 mM MgCl2, 0.3 mM CoA, 0.4 mM ATP and 1 µM OCS (wt), OCS (D) or OCS (K523A, negative control), supplemented with 100 µM oxalate. Reactions were stopped by adding 10 µl of 1 mM HCl and kept on ice. Reaction tubes were centrifuged for 10 minutes at 4 °C, 14,000 rpm on a tabletop centrifuge to remove aggregated proteins and then sequentially injected into a HPLC (Agilent) equipped with a C18 analytical column (Agilent Poroshell 120).

For high-performance liquid chromatography analysis, solvent A was 50 mM ammonium acetate (pH 5.0) and solvent B was 100% methanol. Flow rate was kept at 1 ml/min and at room temperature. Samples were kept at 10 °C in a 96 wells V-bottom plate and sealed to avoid evaporation. The elution method consisted of 0% solvent B for 5 minutes and then a continuous gradient from 0 % to 70 % of solvent B for 15 minutes, and then 100% solvent B for 5 minutes and re-equilibration of the column in solvent A. Samples with only ATP, AMP and CoA were analyzed to define the elution time of each component. AMP production was assessed by measuring the area under the curve between 0.75 and 1.00 minutes of elution. Curves were integrated using GraphPad Prism. All reactions were run in triplicate.

## ACKNOWLEDGMENTS

This project has been supported by FOR 1905 (PerTrans consortium) GA 2519/1-2 to C.G, RA 1781/4-2 to S.R, and 1058/9-1 and 1058/9-2 to M.W from the German Research Foundation (DFG). J. B. has been supported by the Marie Skłodowska-Curie Actions program (grant # 664726) from the European Commission. E.-A. has been supported by the Marie Curie Initial Training Network ‘PERFUME-ITN’ (grant # 316723) from the European Commission. Synchrotron, SAXS and X-ray diffraction data were collected at beamlines P12 and P13 operated by the EMBL Hamburg Unit at the PETRA III storage ring (DESY, Hamburg, Germany) as well as on beamline ID30A-1 at the European Synchrotron Radiation Facility (ESRF), Grenoble, France, respectively. We acknowledge technical support by the SPC facility at EMBL Hamburg Unit. We thank D. Prumbaum and O. Hofnagel (Max Planck Institute of Molecular Physiology, Dortmund) for assistance in cryo-EM data collection.

## Author contributions

JB, EAG, PL, CG and MW designed the project. JB, EAG, PL performed the experiments. JB, PL, CMJ, GC, CG and MW analyzed the data. JB, PL, CG, CMJ, and MW wrote the manuscript. SR, CG and MW supported the project.

## SUPPLEMENT

**Supplementary Table S1:** X-ray data structure determination.

| **X-ray data** | **OCS (wt)** | **OCS (KD)** |
| --- | --- | --- |
| **Accession codes (PDB)** | **8AFF** | **8AFG** |
| Beamline | ID30A-1 (ESRF, Grenoble) | PETRA III P13 (EMBL/DESY, Hamburg) |
| Wavelength (nm) | 0.097 | 0.097 |
| Molecules / asymmetric unit | 12 | 2 |
| Space group | P 2<sub>1</sub> | P 2<sub>1</sub> |
| Cell dimensions |  |  |
| a,b,c (Å) | 109.0, 93.7, 356.5 | 59.9, 125.6, 80.5 |
| $\alpha,\beta,\gamma$ (°) | 90, 93.8, 90 | 90, 96.9, 90 |
| Solvent content (%) | 51 | 42 |
| Resolution range (Å) | 49.20-2.87 (2.97-2.87) | 47.05-2.45 (2.54-2.45) |
| Unique reflections (highest resolution shell) | 162967 (16207) | 43424 (4338) |
| Completeness (%) | 99.2 (99.2) | 99.9 (99.9) |
| I/$\sigma$(I) | 8.6 (1.) | 5.1 (1.1) |
| Multiplicity | 1.9 (1.9) | 4 (4.1) |
| R<sub>merge</sub> (%) | 12.2 (94) | 13.1 (87) |
| CC<sub>1/2</sub> | 0.99 (0.5) | 0.99 (0.51) |
| **Structure refinement** |  |  |
| R-work | 0.230 (0.370) | 0.203 (0.297) |
| R-free | 0.250 (0.371) | 0.237 (0.336) |
| Number of non-hydrogen atoms | 42783 | 7369 |
| Macromolecules | 42466 | 7192 |
| Solvent | 317 | 152 |
| Protein residues | 5395 | 918 |
| R.m.s.d. (bonds) | 0.010 | 0.011 |
| R.m.s.d. (angles) | 1.97 | 2.03 |
| Ramachandran favored (%) | 97.0 | 97.3 |
| Ramachandran allowed (%) | 2.5 | 2.3 |
| Ramachandran outliers (%) | 0.5 | 0.3 |
| Rotamer outliers (%) | 6.3 | 5.2 |
| Clash score | 2.51 | 2.71 |
| Average B-factor | 80.1 | 45.7 |
| Number of TLS groups | 12 | 2 |

**Supplementary Table S2:** Cryo-EM structure determination.

|  |  |
| --- | --- |
| **Accession codes** | **PDB: 8ATD, EMDB: EMD-15646** |
| **Data Collection and processing** |  |
| Microscope | Titan Krios G2 (MPI, Dortmund) |
| Detector | Gatan K2 Summit counting mode |
| Software | EPU ThermoFisher |
| Magnification | 105000x |
| Pixel size (Å/px) | 1.09 |
| Total electron dose (e<sup>-</sup>/Å<sup>2</sup>) | 63 |
| Frames per movie | 60 |
| Defocus range (μm) | -0.7 – 2.2 |
| Micrographs collected | 5593 |
| Micrographs used | 4479 |
| **Reconstruction of the Hexamer** |  |
| Software | TransSPHIRE/ SPHIRE / Relion3.11 |
| Total particles amount | 4000000 |
| Number of particles after 2D classification | 955,374 |
| Number of particles used for refinement | 819,000 |
| Symmetry | C2 |
| Resolution of masked reconstruction at 0.143 Fourier shell correlation (FSC in Å) | 3.1 |
| **Refinement** |  |
| Software | Phenix real space refinement in Phenix |
| Resolution (Å) | 3.1 |
| RMSD bond | 0.01 |
| RMSD angle | 0.971 |
| Model to map fit CC mask | 0.767 |
| **Validation** |  |
| Clash score | 5.12 |
| Ramachandran outliers (%) | 0.12 |
| Ramachandran favored (%) | 96.55 |
| Poor rotamers | 0.22 |
| Molprobrity score | 1.47 |

**Supplementary Table S3:** SAXS data collection.

| **SAMPLE** | **OCS (wt)** | **OCS (wt)** | **OCS K352D** | **OCS K352D** |
| --- | --- | --- | --- | --- |
| **Accession codes (SASBDB)** | SASDPV3 | SASDPW3 | SASDPX3 | SASDPY3 |
| Sample volume | 40 μl | | | |
| Sample conc. (mg/mL) | 2.5 | 0.5 | 2.5 | 0.5 |
| Instrument | PETRA III P12, EMBL/DESY, Hamburg |  |  |  |
| Exposure time (# data frames) | 0.095 s (40) |  |  |  |
| X-ray wavelength | 0.124 nm |  |  |  |
| Sample-to-detector distance | 3.0 m |  |  |  |
| **Primary data analysis** | *PRIMUS* (ATSAS 3.0) |  |  |  |
| Guinier $I(0)$, $\text{cm}^{-1}$ ($\sigma$) | 0.1338<br>(0.0002) | 0.1119<br>(0.0004) | 0.0482<br>(0.0001) | 0.0521<br>(0.0002) |
| $R_g$ (Guinier, nm) ($\sigma$) | 4.67 (0.01) | 4.49 (0.03) | 3.32 (0.01) | 3.31 (0.02) |
| $sR_g$ range/(points used) | 0.42-1.3 (4-71) | 0.49-1.3 (11-75) | 0.44-1.29 (1-93) | 0.3-1.3 (4-112) |
| ***P(r)* analysis** | *GNOM* 5 |  |  |  |
| $I(0)$, POR ($\sigma$) | 0.1339<br>(0.0001) | 0.1124<br>(0.0003) | 0.0484<br>(0.0001) | 0.0522<br>(0.0002) |
| $R_g$ (POR, nm) ($\sigma$) | 4.65 (0.004) | 4.51 (0.02) | 3.36 (0.01) | 3.34 (0.02) |
| $D_{max}$ (nm) | 13 | 14 | 11 | 10 |
| Reciprocal space $p(r)$ fit to data, $\chi^2$ (CorMap P) | 1(0.7) | 1(0.9) | 1(0.3) | 1.1(0.2) |
| Porod volume ($\text{nm}^3$) | 565 | 432 | 163 | 194 |
| ***MW* analysis** |  |  |  |  |
| Calculated MW (kDa) | 60.5 (monomer) |  |  |  |
| MALLS MW (kDa) | 261 (26.48) | 137 (8) | 113 (10) | n.a. |
| SAXS MW (kDa) | 221-373 | 177-264 | 84-96 | 106-127 |
| Hexamer/tetramer (%) | 56/44 | 18/82 | n.a. | n.a. |

### Supplementary Figures

**Figure S1:**
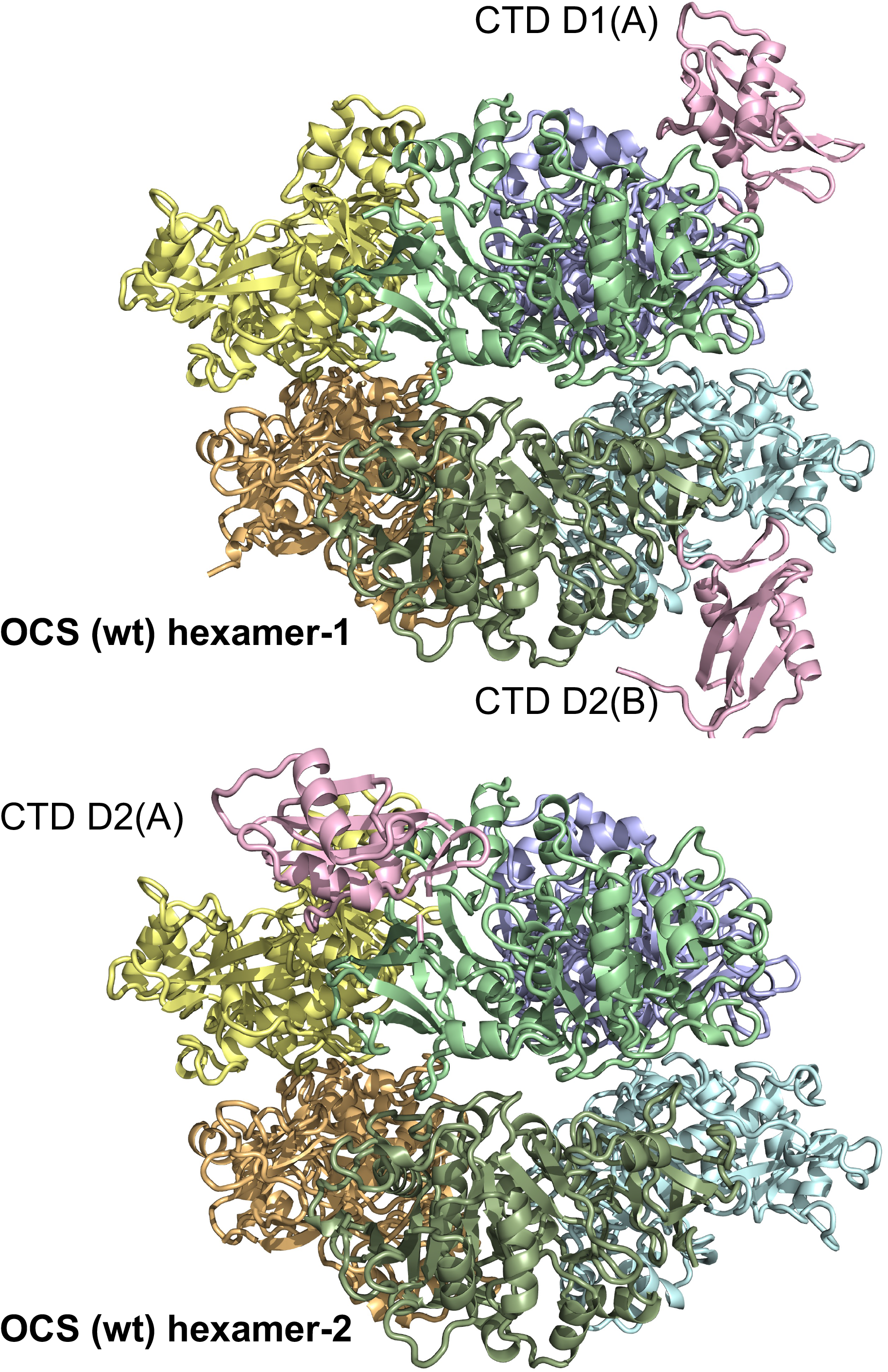
Structural superposition of Hexamer 1 and 2 observed in the asymmetric unit of OCS (wt) crystal. Chain I of hexamer 2 is related to D2(A) in hexamer 1 display a CTD shown in pink, as the CTDs of D1(A) and D2(B).

**Figure S2:**
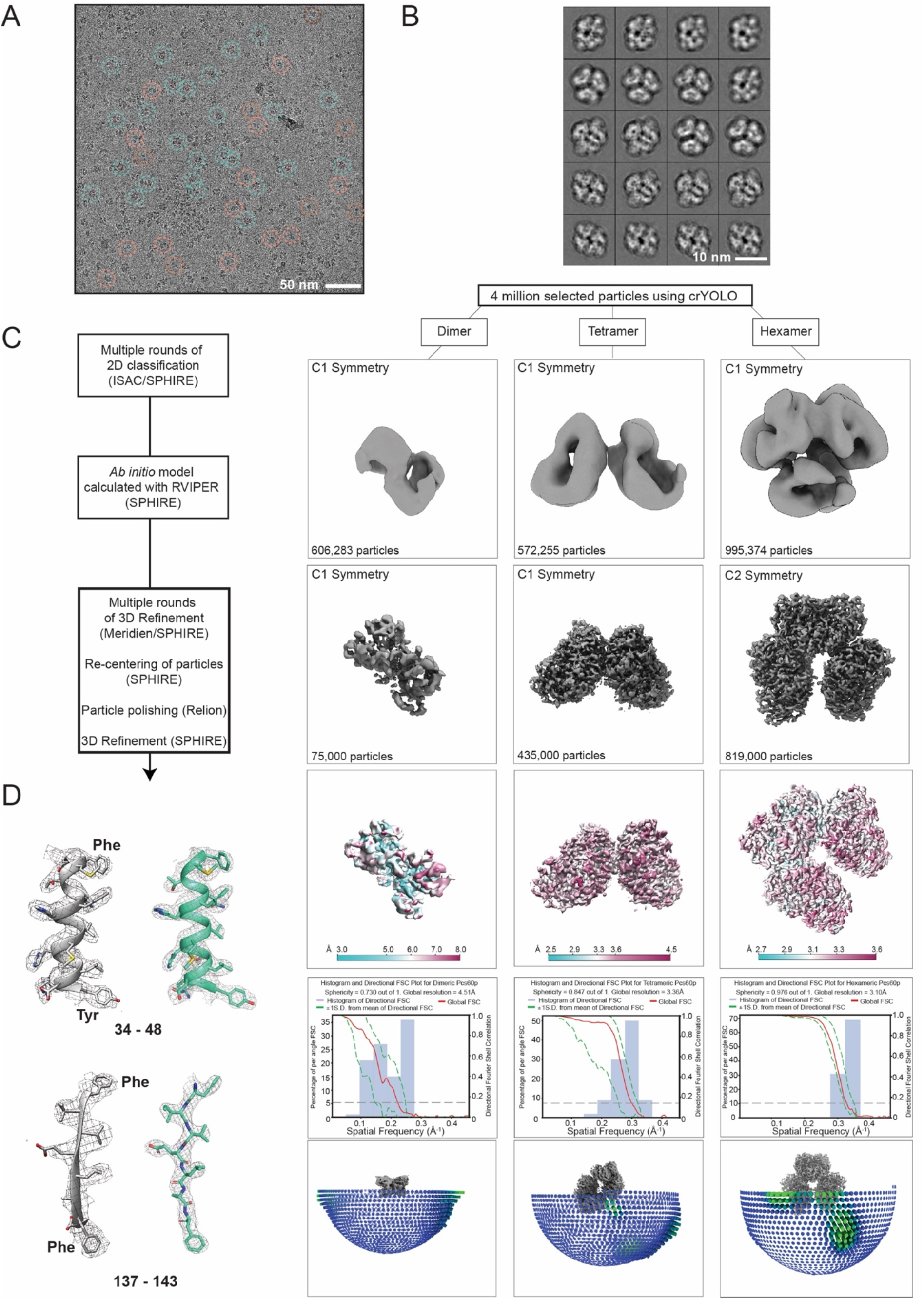
Data processing scheme of OCS cryo-EM structure. **(A)** Representative cryo-EM micrograph of OCS. Particles of different sizes are highlighted by circles of different colors. **(B)** Representative 2D classes of OCS. **(C)** Flow chart of the cryo-EM data-processing procedure to generate the maps of the dimer, tetramer and hexamer. Local resolution and 3D FSC curves are shown for each OCS oligomer. **(D)** Comparison of OCS cryo-EM electron density map and refined model (in grey) with OCS crystal structure map and refined model (in cyan green).

**Figure S3:**
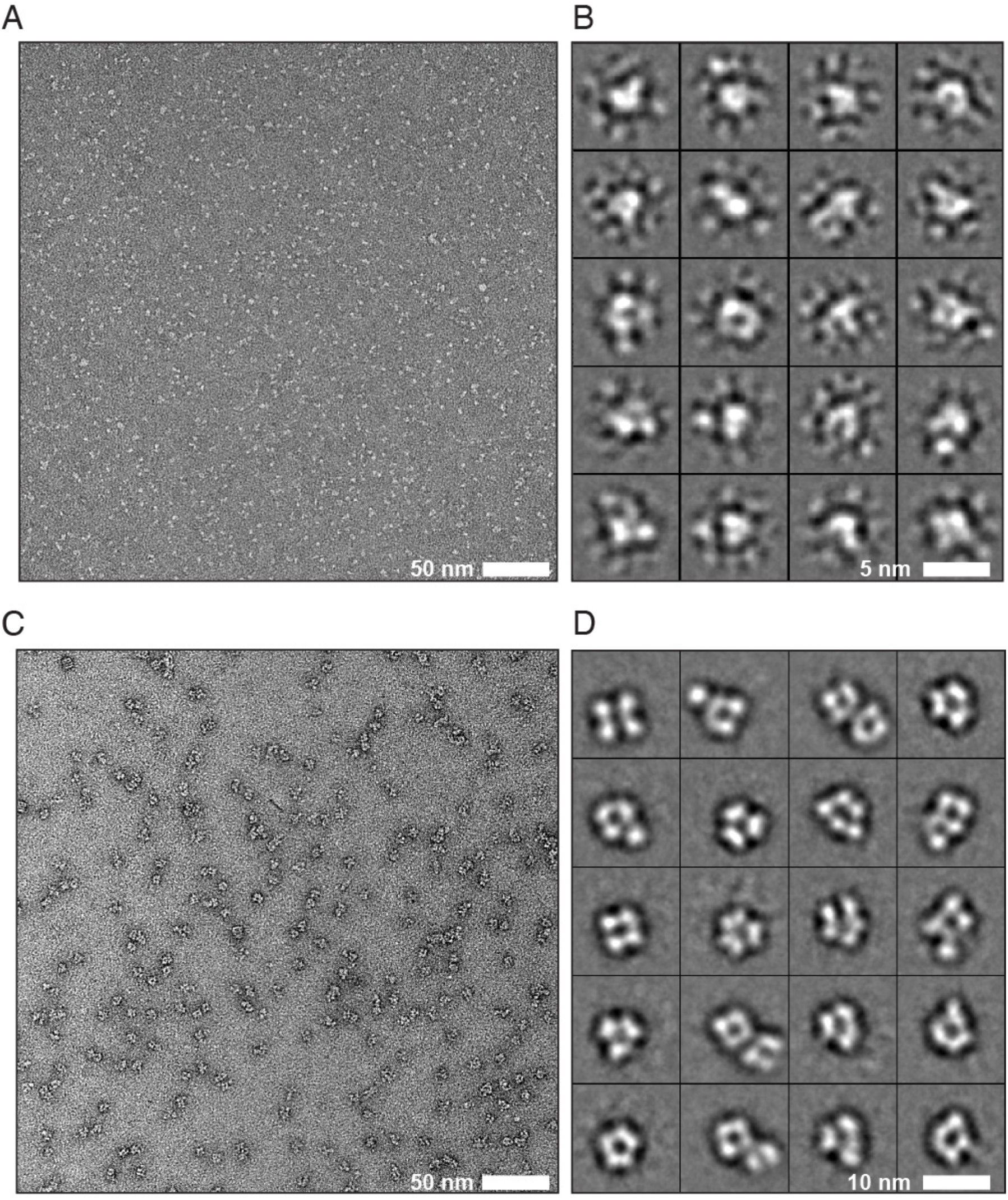
Negative stain EM analysis of OCS (wt). **(A)** Representative negative stain electron micrograph of OCS (wt). **(B)** 2D classes show particles in different orientations. Small particles are not aligning well, but the majority of the class averages is consistent with the size of dimeric OCS. **(C)** Representative micrograph of crosslinked OCS (wt). **(D)** Reference-free 2D classes of crosslinked OCS (wt), showing the characteristic views of the hexameric assembly. Scale bars indicate the dimensions of particles.

**Figure S4:**
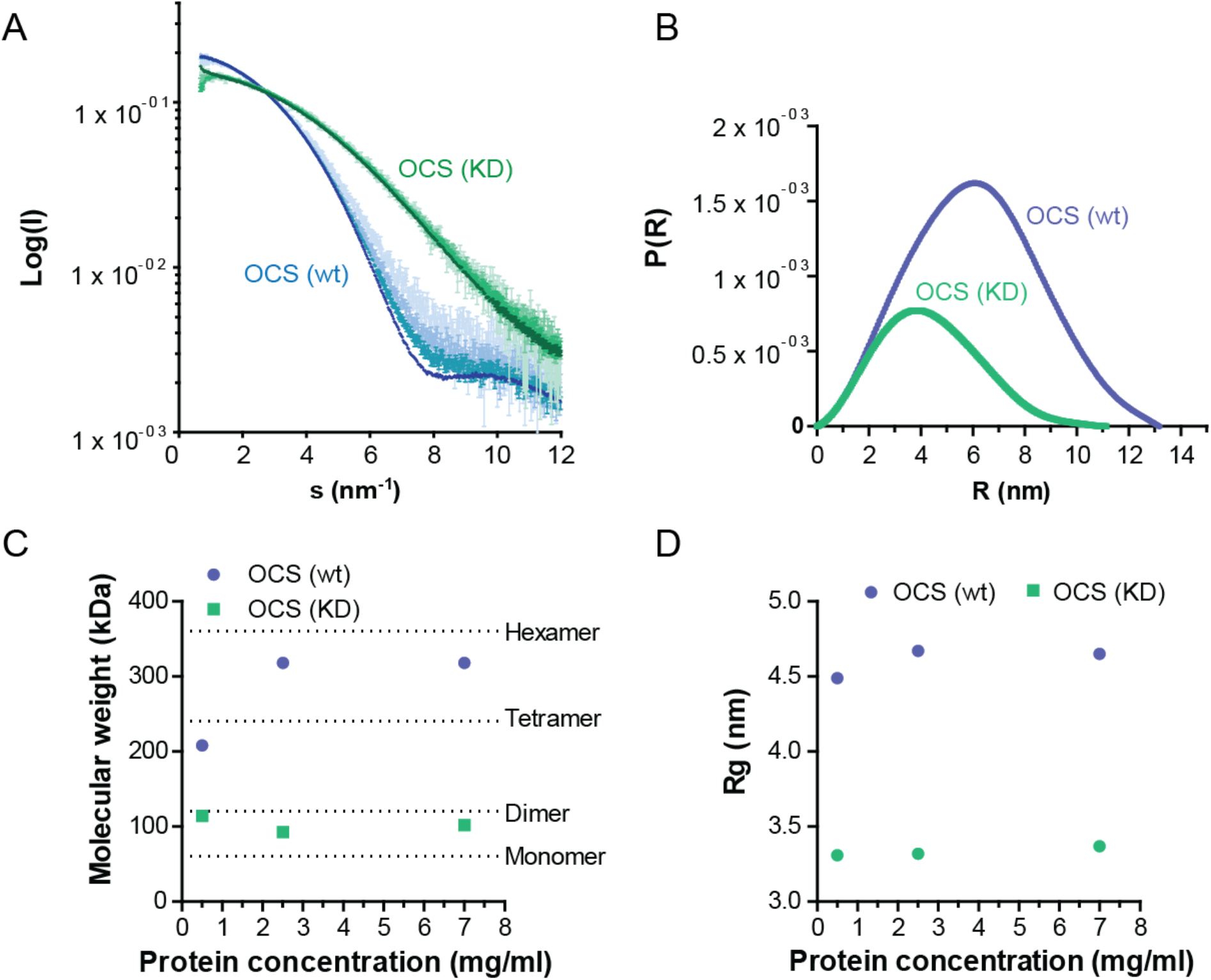
**(A)** SAXS profiles of OCS (wt) in different shades of blue spanning 7 mg/mL (dark blue) through to 0.5 mg/mL (light blue) and the corresponding SAXS data measured from the OCS (KD) mutant spanning the same concentration range (from dark to light green). To more clearly demonstrate the concentration-dependent differences in the SAXS data for the wt OCS, and especially differences in the mid- to higher *s*-range (0.4–1.2 nm^-1^), the datasets have been scaled to the same *j*(0). Similar scaling to a common *I*(0) for the KD mutant was also applied, showing minimal concentration dependence in the data across the concentration series. **(B)** The *p*(*r*) plot of OCS (wt, blue) and OCS (KD, green) at a concentration of 2.5 mg/ml. **(C and D)** Estimated MW (C) and *R_g_* (D) of OCS (wt, blue) and OCS (KD, green) at different protein concentrations.

**Figure S5:**
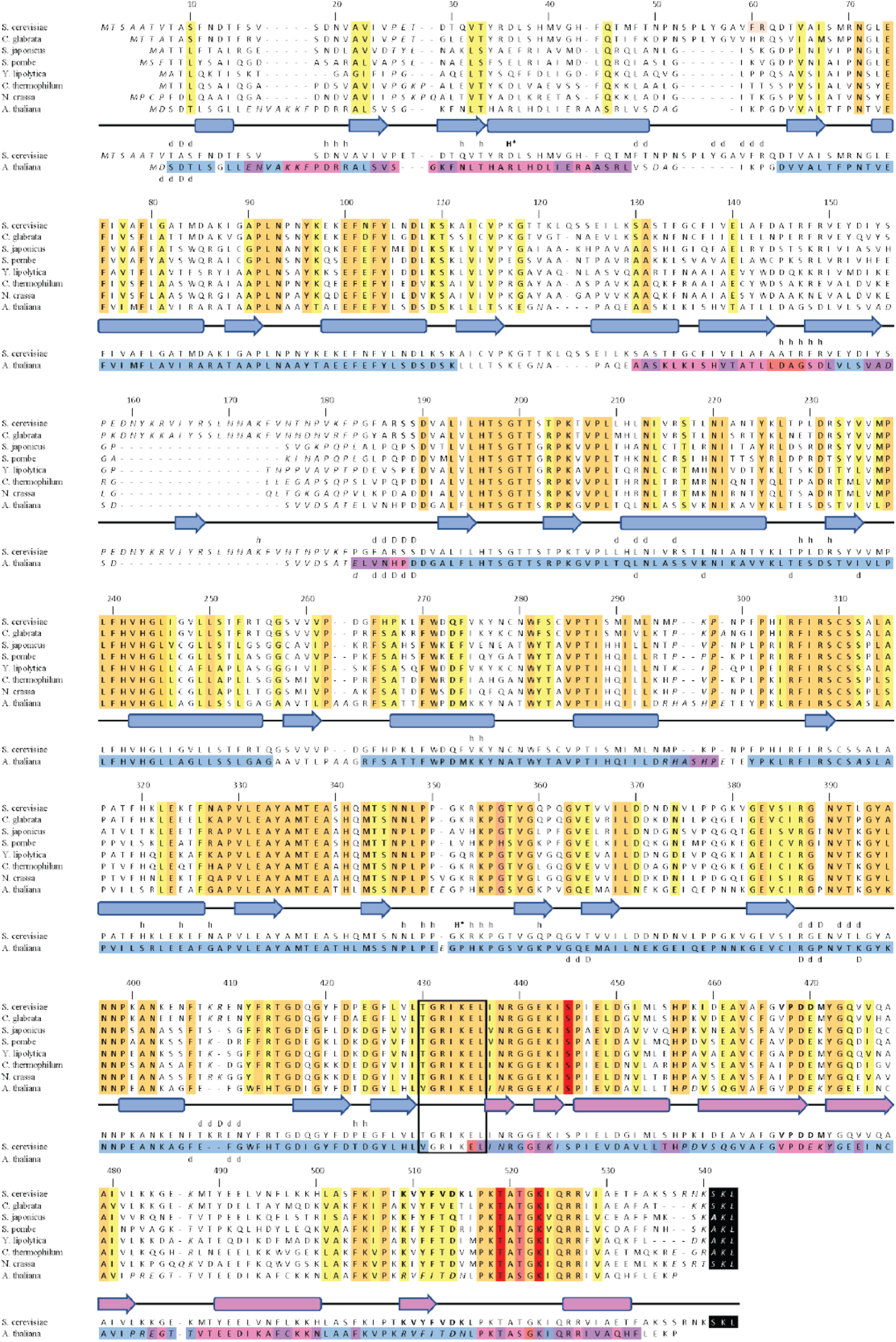
**(A)** Sequence alignment of fungal OCS from *S. cerevisiae, Candida glabrata, Schizosaccharomyces japonicus and pombe, Yarrowia lipolytica, Chaetomium thermophilum, Neurospora crassa* and plant OCS from *A. thaliana.* (A and B). Residue numbers relate to the *S. cerevisiae* OCS sequence. Conserved and invariant residues positions are highlighted in yellow and orange, respectively. Conserved residues that contribute to specific NTD/CTD arrangements are highlighted in red. The C-terminal PTS1 motifs of the fungal OCS sequences are highlighted in black. The NTD/CTD hinge segment is boxed. Structure-based sequence alignment for OCS with known structures from *S. cerevisiae* and *A. thaliana* is shown below. Residue positions where the alignment is uncertain are in italics. Residue positions involved in dimer interfaces are indicated by “d”. Residue positions that are involved in hexameric interfaces by dimer self-assembly are indicated by “h”. Those residues that contribute to specific side-chain mediated interface interactions are indicated in capital letters. Secondary structural elements corresponding to the *S. cerevisiae* OCS structure are shown as well. The rainbow colors of the *A. thaliana* OCS sequence correspond to the rmsd values of the superimposed OCS structures from *S. cerevisiae* and A. thaliana **(cf. Supplementary Figure S6).**

**Figure S6:**
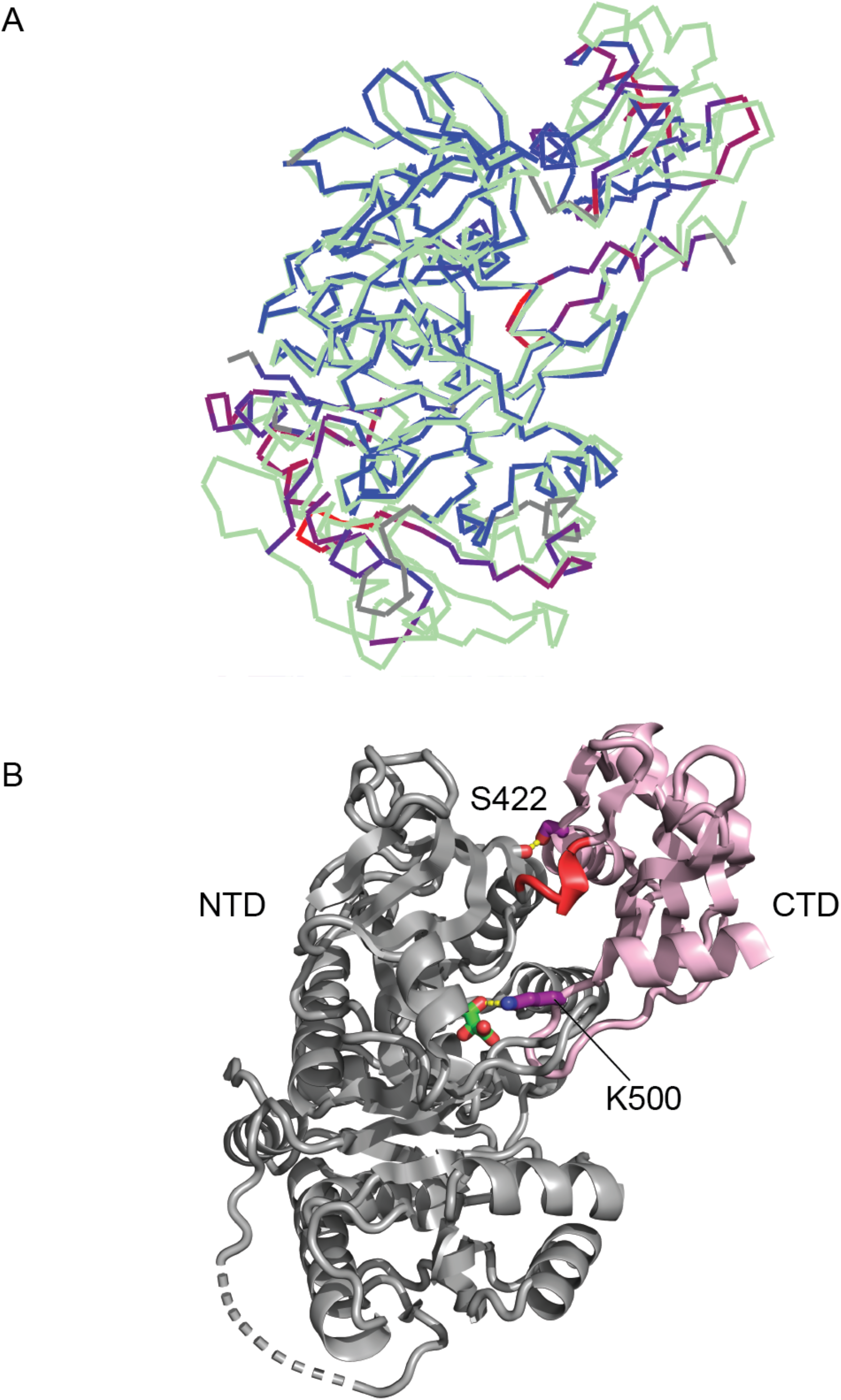
**(A)** Structural alignment of a *S. cerevisiae* OCS (wt) protomer (pale green) and *A. thaliana* OCS protomer (orange, PDB: 5IE0), shown in rainbow colors depending on rmsd, ranging from blue (minimum distance) to red (maximum distance). Sequence segments without a match are in grey. The NTDs and CTDs of both structures were superimposed separately, resulting in rmsd of 0,86 Å (1749 matching atoms) and 1,27 Å (632 matching atoms), respectively. The ALIGN option in PyMol was used for superposition. Coloring was done with the COLORBYRMSD plugin in PyMol. **(B)** Cartoon representation of the NTD/CTD arrangement of OCS from *A. thaliana.* The key interaction between S422 from the CTD and a main chain carbonyl group from the NTD is conserved in the OCS structures from *S. cerevisiae* and *A. thaliana* (***cf*.** Figure 2D**, left panel)**. The *A. thaliana* OCS structure (5ie3) contains a tartrate ion in the active site, which pulls K500 from the CTD active site loop towards the NTD by an additional specific interaction.

**Figure S7:**
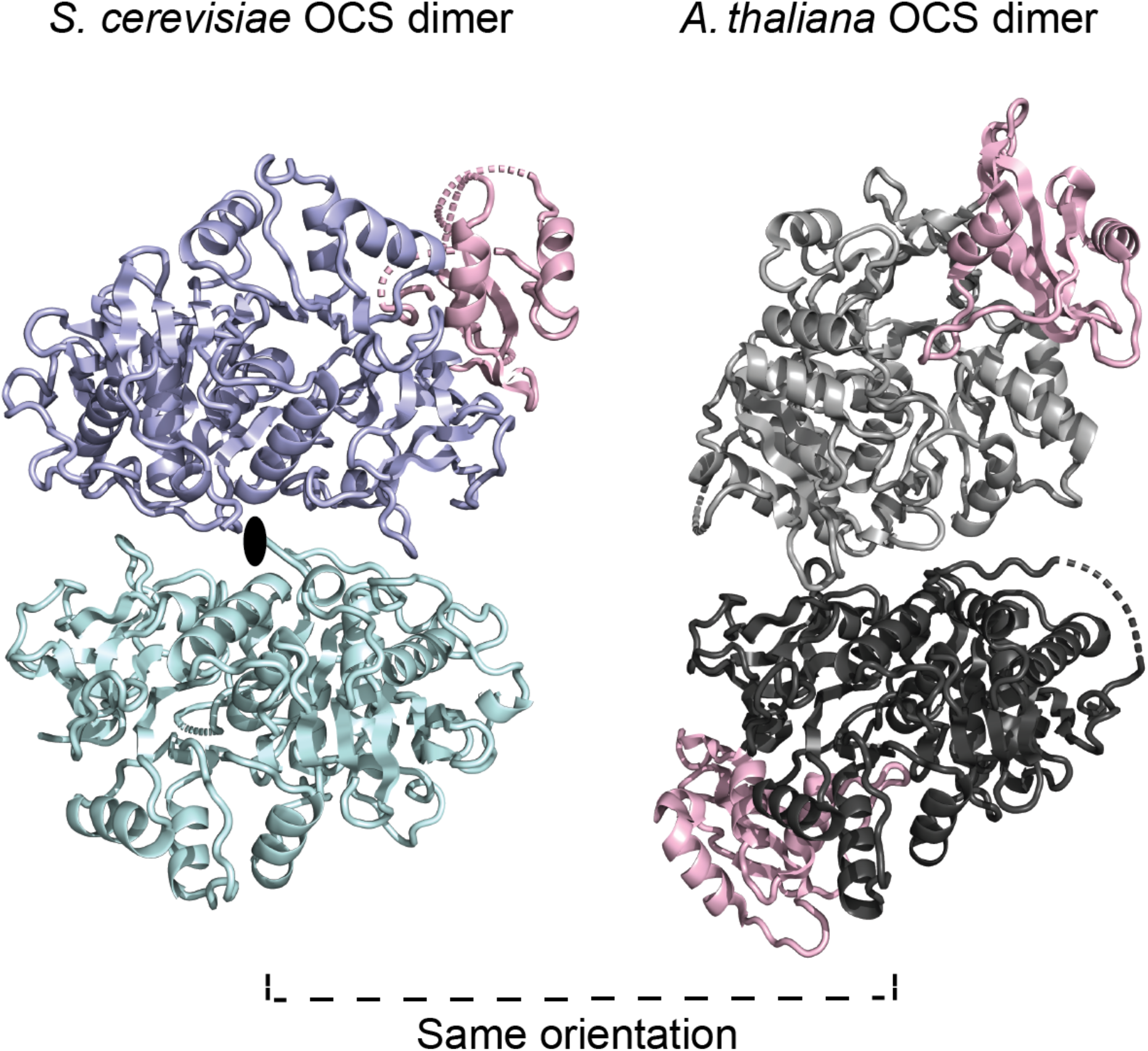
Non-conserved dimer arrangement in *S. cerevisiae* and *A. thaliana* OCS. The lower subunits are in identical orientation. The dimer axis of the *S. cerevisiae* OCS is vertical to the document plane and is indicated. Colors for *S. cerevisiae* OCS are as in Figures 1 and 2. The NTDs of the *A. thaliana* OCS structure are shown in different grey shades.

## Notes

### Competing Interest Statement

The authors have declared no competing interest.

## REFERENCES

Adams, Paul D., Pavel V. Afonine, Gábor Bunkóczi, Vincent B. Chen, Nathaniel Echols, Jeffrey J. Headd, Li Wei Hung, et al. 2011. “The Phenix Software for Automated Determination of Macromolecular Structures.” Methods. https://doi.org/10.1016/j.ymeth.2011.07.005.

Blanchet, Clement E., Alessandro Spilotros, Frank Schwemmer, Melissa A. Graewert, Alexey Kikhney, Cy M. Jeffries, Daniel Franke, et al. 2015. “Versatile Sample Environments and Automation for Biological Solution X-Ray Scattering Experiments at the P12 Beamline (PETRA III, DESY).” Journal of Applied Crystallography. https://doi.org/10.1107/S160057671500254X.

Blobel, Fabian. 1996. “Identification of a Yeast Peroxisomal Member of the Family of AMP-Binding Proteins.” European Journal of Biochemistry 240 (2): 468–76. https://doi.org/10.1111/j.1432-1033.1996.0468h.x.

DeLano, W L. 2020. “The PyMOL Molecular Graphics System, Version 2.3.” Schrödinger LLC.

Emsley, P., B. Lohkamp, W. G. Scott, and K. Cowtan. 2010. “Features and Development of Coot.” Acta Crystallographica Section D: Biological Crystallography. https://doi.org/10.1107/S0907444910007493.

Fan, Minrui, Yang Xiao, Mei Li, and Wenrui Chang. 2016. “Crystal Structures of Arabidopsis Thaliana Oxalyl-CoA Synthetase Essential for Oxalate Degradation.” Molecular Plant 9 (9): 1349–52. https://doi.org/10.1016/j.molp.2016.06.002.

Foster, Justin, and Paul A. Nakata. 2014. “An Oxalyl-CoA Synthetase Is Important for Oxalate Metabolism in Saccharomyces Cerevisiae.” FEBS Letters 588 (1): 160–66. https://doi.org/10.1016/j.febslet.2013.11.026.

Franke, D., M. V. Petoukhov, P. V. Konarev, A. Panjkovich, A. Tuukkanen, H. D. T. Mertens, A. G. Kikhney, et al. 2017. “*ATSAS 2.8* : A Comprehensive Data Analysis Suite for Small-Angle Scattering from Macromolecular Solutions.” Journal of Applied Crystallography 50 (4): 1212–25. https://doi.org/10.1107/S1600576717007786.

Franke, Daniel, Alexey G. Kikhney, and Dmitri I. Svergun. 2012. “Automated Acquisition and Analysis of Small Angle X-Ray Scattering Data.” Nuclear Instruments and Methods in Physics Research, Section A: Accelerators, Spectrometers, Detectors and Associated Equipment. https://doi.org/10.1016/j.nima.2012.06.008.

Goodsell, David S., and Arthur J. Olson. 2000. “Structural Symmetry and Protein Function.” Annual Review of Biophysics and Biomolecular Structure. https://doi.org/10.1146/annurev.biophys.29.1.105.

Gulick, Andrew M., Vincent J. Starai, Alexander R. Horswill, Kristen M. Homick, and Jorge C. Escalante-Semerena. 2003. “The 1.75 Å Crystal Structure of Acetyl-CoA Synthetase Bound to Adenosine-5′-Propylphosphate and Coenzyme A.” Biochemistry. https://doi.org/10.1021/bi0271603.

Gulick, Andrew M, W D Mcelroy, M Deluca, and J Travis. 2009. “Conformational Dynamics in the Acyl-CoA Synthetases, Adenylation Domains of Non-Ribosomal Peptide Synthetases, and Firefly Luciferase” 4 (10): 811–27. https://doi.org/10.1021/cb900156h.

Hagen, Stefanie, Friedel Drepper, Sven Fischer, Krisztian Fodor, Daniel Passon, Harald W. Platta, Michael Zenn, et al. 2015. “Structural Insights into Cargo Recognition by the Yeast PTS1 Receptor *.” Journal of Biological Chemistry 290 (44): 26610–26. https://doi.org/10.1074/jbc.M115.657973.

Hayward, Steven, and Richard A. Lee. 2002. “Improvements in the Analysis of Domain Motions in Proteins from Conformational Change: DynDom Version 1.50.” Journal of Molecular Graphics and Modelling. https://doi.org/10.1016/S1093-3263(02)00140-7.

Jumper, John, Richard Evans, Alexander Pritzel, Tim Green, Michael Figurnov, Olaf Ronneberger, Kathryn Tunyasuvunakool, et al. 2021. “Highly Accurate Protein Structure Prediction with AlphaFold.” Nature. https://doi.org/10.1038/s41586-021-03819-2.

Kabsch, Wolfgang. 2010. “Integration, Scaling, Space-Group Assignment and Post-Refinement.” Acta Crystallographica Section D: Biological Crystallography. https://doi.org/10.1107/S0907444909047374.

Kikhney, Alexey G., Clemente R. Borges, Dmitry S. Molodenskiy, Cy M. Jeffries, and Dmitri I. Svergun. 2020. “SASBDB: Towards an Automatically Curated and Validated Repository for Biological Scattering Data.” Protein Science. https://doi.org/10.1002/pro.3731.

Konarev, Petr V., Vladimir V. Volkov, Anna V. Sokolova, Michel H.J. Koch, and Dmitri I. Svergun. 2003. “PRIMUS: A Windows PC-Based System for Small-Angle Scattering Data Analysis.” Journal of Applied Crystallography. https://doi.org/10.1107/S0021889803012779.

Krissinel, Evgeny. 2012. “Enhanced Fold Recognition Using Efficient Short Fragment Clustering.” Journal of Molecular Biochemistry.

Krissinel, Evgeny, and Kim Henrick. 2007. “Inference of Macromolecular Assemblies from Crystalline State.” Journal of Molecular Biology. https://doi.org/10.1016/j.jmb.2007.05.022.

Lee, Richard A., Moe Razaz, and Steven Hayward. 2003. “The DynDom Database of Protein Domain Motions.” Bioinformatics. https://doi.org/10.1093/bioinformatics/btg137.

Liebschner, Dorothee, Pavel V. Afonine, Matthew L. Baker, Gábor Bunkoczi, Vincent B. Chen, Tristan I. Croll, Bradley Hintze, et al. 2019. “Macromolecular Structure Determination Using X-Rays, Neutrons and Electrons: Recent Developments in Phenix.” Acta Crystallographica Section D: Structural Biology. https://doi.org/10.1107/S2059798319011471.

Maksay, Gábor, and Joseph A. Marsh. 2017. “Signalling Assemblies: The Odds of Symmetry.” Biochemical Society Transactions. https://doi.org/10.1042/BST20170009.

Marsh, Joseph A., and Sarah A. Teichmann. 2015. “Structure, Dynamics, Assembly, and Evolution of Protein Complexes.” Annual Review of Biochemistry. https://doi.org/10.1146/annurev-biochem-060614-034142.

McCoy, Airlie J., Ralf W. Grosse-Kunstleve, Paul D. Adams, Martyn D. Winn, Laurent C. Storoni, and Randy J. Read. 2007. “Phaser Crystallographic Software.” Journal of Applied Crystallography. https://doi.org/10.1107/S0021889807021206.

Moriya, Toshio, Michael Saur, Markus Stabrin, Felipe Merino, Horatiu Voicu, Zhong Huang, Pawel A. Penczek, Stefan Raunser, and Christos Gatsogiannis. 2017. “High-Resolution Single Particle Analysis from Electron Cryo-Microscopy Images Using SPHIRE.” Journal of Visualized Experiments. https://doi.org/10.3791/55448.

Murshudov, Garib N., Pavol Skubák, Andrey A. Lebedev, Navraj S. Pannu, Roberto A. Steiner, Robert A. Nicholls, Martyn D. Winn, Fei Long, and Alexei A. Vagin. 2011. “REFMAC5 for the Refinement of Macromolecular Crystal Structures.” Acta Crystallographica Section D: Biological Crystallography. https://doi.org/10.1107/S0907444911001314.

Pettersen, Eric F., Thomas D. Goddard, Conrad C. Huang, Gregory S. Couch, Daniel M. Greenblatt, Elaine C. Meng, and Thomas E. Ferrin. 2004. “UCSF Chimera - A Visualization System for Exploratory Research and Analysis.” Journal of Computational Chemistry. https://doi.org/10.1002/jcc.20084.

Pettersen, Eric F., Thomas D. Goddard, Conrad C. Huang, Elaine C. Meng, Gregory S. Couch, Tristan I. Croll, John H. Morris, and Thomas E. Ferrin. 2021. “UCSF ChimeraX: Structure Visualization for Researchers, Educators, and Developers.” Protein Science. https://doi.org/10.1002/pro.3943.

Stabrin, Markus, Fabian Schoenfeld, Thorsten Wagner, Sabrina Pospich, Christos Gatsogiannis, and Stefan Raunser. 2020. “TranSPHIRE: Automated and Feedback-Optimized on-the-Fly Processing for Cryo-EM.” Nature Communications. https://doi.org/10.1038/s41467-020-19513-2.

Studier, F William. 2005. “Protein Production by Auto-Induction in High Density Shaking Cultures.” Protein Expression and Purification.

Svergun, D., C. Barberato, and M. H. Koch. 1995. “CRYSOL - A Program to Evaluate X-Ray Solution Scattering of Biological Macromolecules from Atomic Coordinates.” Journal of Applied Crystallography. https://doi.org/10.1107/S0021889895007047.

Veevers, Ruth, and Steven Hayward. 2019. “Methodological Improvements for the Analysis of Domain Movements in Large Biomolecular Complexes.” Biophysics and Physicobiology. https://doi.org/10.2142/biophysico.16.0_328.

Wagner, Thorsten, Felipe Merino, Markus Stabrin, Toshio Moriya, Claudia Antoni, Amir Apelbaum, Philine Hagel, et al. 2019. “SPHIRE-CrYOLO Is a Fast and Accurate Fully Automated Particle Picker for Cryo-EM.” Communications Biology. https://doi.org/10.1038/s42003-019-0437-z.

Williams, Hibbard E., and Lloyd H. Smith. 1968. “Disorders of Oxalate Metabolism.” The American Journal of Medicine. https://doi.org/10.1016/0002-9343(68)90207-6.

Yang, Zhengfan, Jia Fang, Johnathan Chittuluru, Francisco J. Asturias, and Pawel A. Penczek. 2012. “Iterative Stable Alignment and Clustering of 2D Transmission Electron Microscope Images.” Structure. https://doi.org/10.1016/j.str.2011.12.007.

Zhang, Kai. 2016. “Gctf: Real-Time CTF Determination and Correction.” Journal of Structural Biology. https://doi.org/10.1016/j.jsb.2015.11.003.

Zheng, Shawn Q., Eugene Palovcak, Jean Paul Armache, Kliment A. Verba, Yifan Cheng, and David A. Agard. 2017. “MotionCor2: Anisotropic Correction of Beam-Induced Motion for Improved Cryo-Electron Microscopy.” Nature Methods. https://doi.org/10.1038/nmeth.4193.

Zivanov, Jasenko, Takanori Nakane, and Sjors H.W. Scheres. 2019. “A Bayesian Approach to Beam-Induced Motion Correction in Cryo-EM Single-Particle Analysis.” IUCrJ. https://doi.org/10.1107/S205225251801463X.

